# LC-MS analysis of RNA through hydrogen bonding of alkylamines

**DOI:** 10.1101/2025.03.18.643832

**Authors:** Robert L. Ross, Michael J. Sutton, Keeley Murphy, Roberto Gamez, Min Du, Michael G. Bartlett

## Abstract

Nucleic acid oligonucleotides represent a promising therapy for treating disease. Production of these biopolymers happens synthetically which can produce impurities concomitant with the desired product. The use of liquid chromatography coupled to high resolution mass spectrometry is a robust analytical technique for determination of sample purity. Traditionally, ion pairing reagents are used to facilitate chromatography separation before sampling with the mass spectrometer. The choice of reagent for analysis can impact the analysis as the length of the oligonucleotide increases. This is in part due to the thermodynamics of transitioning the oligonucleotide from the liquid phase to the gas phase. Here we show that the use of a hydrogen bond donating alkylamine assists with overcoming the energy barrier of phase transition by bonding to the nucleobase as well as the phosphodiester backbone. By increasing the relative hydrophobicity of the oligonucleotide, the transition to the gas phase is increased producing a cleaner mass spectrum. Furthermore, we show that by using a hydrogen bond donating alkylamine, we can remove the fluoroalcohol reagent commonly used in ion pairing mass spectrometry of nucleic acids.

## Introduction

The use of RNA based therapeutics^1,2^ continues to grow, and with it the need for analytical methods to ensure quality and efficacy of these synthetic drugs.^3^ The introduction of CRISPR^4^ gene editing based therapies utilizes longer RNA sequences than that of antisense^5^ or interfering RNA^6^ therapies. In CRISPR/Cas13^7,8^ systems specific guide RNAs (sgRNA)^9^ have lengths of ∼ 40 nucleotides (nt) where the Cas9 system utilizes lengths of 100 nts or more and can carry multiple modified nucleotides.^10,11^ These oligonucleotides present challenges analytically,^12^ as they are produced synthetically^13^ one nucleotide at a time^14^ and generate considerable impurities, “failure sequences” with each monomer addition.

The use of ion pair liquid chromatography hyphenated to high resolution accurate mass spectrometry has provided a solid analytical platform in oligonucleotide analysis and has been effectively used to retain and separate complex mixtures for decades^15-17^ The main retention mechanism for this separation has involved intercalation of the alkylamine into a hydrophobic stationary phase presenting a surface positive charge allowing ionic interaction with the phosphate backbone of the oligonucleotide^18^ It has also been shown that the alkylamine in the mobile phase will directly interact with the phosphate backbone masking the negative change and thereby increasing the hydrophobicity of the oligonucleotide^19^.

The alkylamine / fluoroalcohol mobile phase system has repeatedly been shown to be a complex mixture of competing chemistries that influence both chromatographic retention and electrospray desorption. Chromatographic retention follows the on-off model, which has been well described by Guillarme et al.^20^ The on/off mechanism is primarily driven by alkylamine hydrophobicity, alkylamine concentration, column temperature, counterion type and mobile phase pH. One of the most notable features of this mechanism is the lack of influence of sequence or composition of the oligonucleotide.^21^ This means that all oligonucleotides of the same length will elute together. This feature makes impurity analysis more dependent on the use of electrospray ionization mass spectrometry to distinguish impurities by their differing molecular weights.

Three models explain oligonucleotide ionization during the electrospray process. In the Charge Residue model,^22^ an electrosprayed droplet will undergo multiple fission events^23^ due to coulombic repulsion of the surface charges, until a single molecule is left in the droplet. The residual surface charges are then transferred to the molecule upon evaporation of the droplet. In the Ion Desorption model,^24^ the oligonucleotide overcomes the thermodynamic energy barrier to desorb from the droplet with help from the electric field. In this model the ionic interaction between alkylamine and phosphate dissociates due to an increasing pH as the electrosprayed droplet evaporates. Once the droplet’s pH increases to the alkylamine’s pKa, the ionic interaction is interrupted, and the electronegative backbone of the oligonucleotide is then acted upon by the pervasive electric field to transition from the liquid to the gas phase. A third model, the Chain Ejection model,^25^ where a biopolymer escapes from the droplet, gradually by the subunit in a linear fashion. The transition from the liquid phase to the gas phase is thermodynamically unfavorable for a biopolymer. For the latter two models, the energy barrier is overcome with assistance from the electric field, the analytes innate hydrophobicity, and adduction.^26^

There are many other critical features of the alkylamine / fluoroalcohol mobile phase system. Among these important features that have been noted during method development for ion-pair reversed phase LC-MS based separations include alkylamine Henry’s Law constant,^27^ fluoroalcohol hydrophobicity,^28^ alkylamine oxidation,^29^ alkylamine aggregation^30^ and pH dependent non-specific adsorption. It is, therefore, important to understand that method development is often a careful balancing act among the various attributes and conditions of the reagents involved.

However, what is often less appreciated is that ion-pair chromatography with non-alkane based stationary phases have shown the ability to interact with both faces of the oligonucleotide. This has allowed these stationary phases to distinguish between different compositions and sequences of oligonucleotides. This view is supported by studies involving phenyl^31^ and cholesterol^32^ columns which, in addition to ionic interactions mediated by alkylamines, also demonstrated complex pi-stacking interactions with the nucleobases. An additional new observation has been made in hydrophilic interaction chromatography, which has noted that acceptor-donor interactions between the nucleobases and the stationary phase are also significant contributors to retention.^33^ However, until this study these types of interactions have not been noted in ion-pair reversed-phase chromatography. Here we show that the alkylamines used in ion-pairing reverse phase chromatography of nucleic acids can form a hydrogen bond with the nucleobase. The added hydrophobicity from the addition affects chromatographic retention and enhances electrospray ionization during LC-MS sampling. The increased signal allows for LC-MS sampling of nucleic acids without the need for HFIP.

## Experimental

### Reagents

N, N-dibutylamine, triethylamine, diethylamine, Optima Grade UHPLC Water and Acetonitrile, LC-MS Grade ammonium acetate, acetic acid and DNA/RNase free water were purchased from Thermo Fisher Scientific.

### Samples

Non-modified RNA 40mer, RNA 100mer, polydeoxyadenosine (15mer) and polydeoxyinosine (15mer) were purchased through Integrated DNA Technology.

40mer RNA Sequence AUGGCCCCCAAGAAGAAGCGGAAGGUGGGCAUCCACGGCG

100mer RNA Sequence

CGUGCCCGACACCAAGGUGAACUUCUACGCCUGGAAGCGGAUGGAGGUGGGCCAGCAGGCCGUGGAGGUGUGGCAGGGCCUGGCCCUGCUGAGCGAGGCC

### Liquid Chromatography Mass Spectrometry

All separations were accomplished on a Thermo Scientific™ Vanquish™ Horizon Binary UHPLC system (Thermo Fisher Scientific) using a DNAPac™ RP column, Accucore™ C8 HPLC Columns, or Javelin C4 column (Thermo Fisher Scientific). Mobile phase A consisted of 5mM dibutyl amine (Thermo Fisher Scientific), 25mM hexafluoro-2-propanol (Thermo Fisher Scientific). Mobile phase B consisted of 100% acetonitrile. An elution gradient of 5% B (from 0 to 0.5 min), 40% B at 7.0 min, 90% B at 7.1 held for 0.5 min, returning to 10% B at 7.51 min at a flow rate of 400 µL min^-1^ was used. The column temperature was set at 75 °C for DNAPac and 35°C for C8 and C4 columns. High resolution accurate mass analysis was performed on an Thermo Scientific™ Orbitrap Ascend™ BioPharma Tribrid™ mass spectrometer (Thermo Fisher Scientific) interfaced with a heated electrospray (H-ESI) source.

### Intact Mass Analysis

Full scan acquisition was acquired in negative polarity using Intact Protein mode with low pressure setting, with a resolution of either 480,000 or 240,000 at 400 m/z, mass range 800-3000 m/z, automatic gain control (AGC) 4e5, and max injection time (IT) 500 ms; quadrupole isolation of 1 m/z; and radio frequency (RF) of 50%; Global instrument conditions were sheath gas, auxiliary gas, and sweep gas of 50, 10 and 1 arbitrary units, respectively; ion transfer tube temperature of 350 °C; vaporizer temperature of 320 °C; and spray voltage of 3500 V. Data was acquired using Xcalibur™ 4.7 and analyzed with Freestyle™ 1.8 and BioPharma Finder™ 5.3.

Deconvolution was performed using Thermo Scientific BioPharma Finder™ software 5.3, Intact mass analysis workflow, using the Xtract deconvolution algorithm. The chromatogram trace type was TIC, and m/z range of 800 to 3000. Source spectra method was averaged over selected time. The deconvolution algorithm was Xtract with an output mass range dependent on the constructs size with a S/N threshold of 3 and a relative abundance threshold (%) of 1. Charge range was set from 10 to 35 with a minimum number of detected charges set to5 and using the sequence specific table. Polarity was set to negative charge. Under identification, sequence matching mass tolerance was set to 10 ppm with multi-consensus component merge mass tolerance at 10 ppm.

## Results & Discussion

### DBA and ACN adducts at lower charge states

Examining the chemical structure of a nucleic acid shows that the number of hydrogen bond acceptor sites outnumber donor sites by 2:1 (**Figure 1**). Hydrogen bonding between the nucleobases results in secondary and tertiary structure of nucleic acids, as well as RNA:Protein complexes, therefore the ability of a ssRNA molecule to form a hydrogen bond with a chemical element in solution is highly probable.^34^ Therefore, we sought to explore the use of the secondary amine, dibutylamine (DBA) to exploit the hydrogen bonding sites on the nucleobases to increase the overall hydrophobicity of the RNA strand with the goal of enhancing the electrospray signal.^35^ Various amines were initially tried; however, we chose DBA based on its solubility and overall alkyl chain length. Furthermore, the use of acetonitrile as the organic phase was due to the aprotic solvent’s eluotropic strength and boiling point. Secondly, the polystyrene divinylbenzene (PS-DVB) stationary phase is resistant to solubility issues of hexafluoro isopropanol with acetonitrile reported with silica based C18 materials,^36^ at least at the concentrations used in this study (< 100 mM).

**Figure 1:**
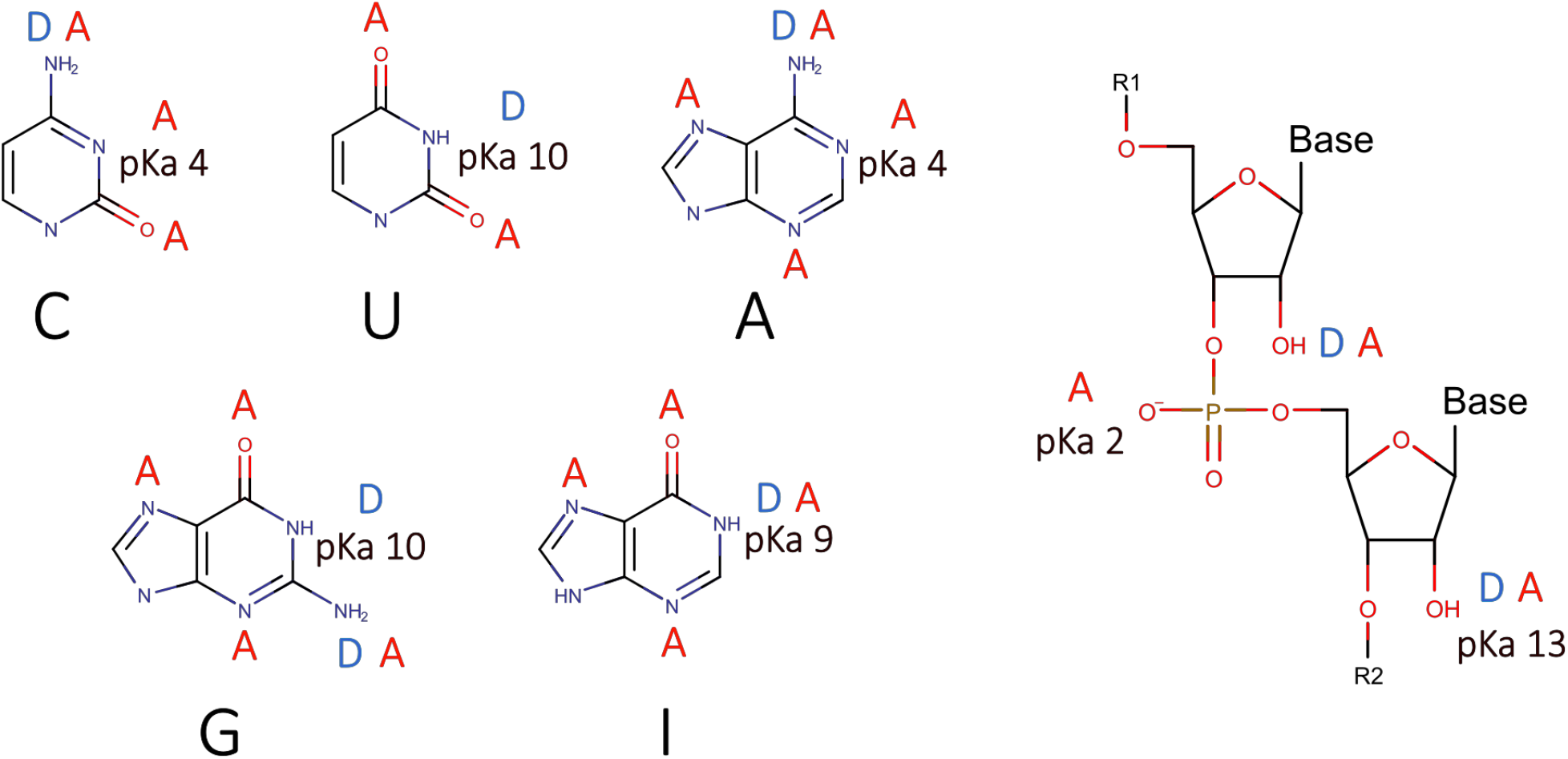
Cartoon showing the availability of hydrogen bonding sites as well as pKa values for the canonical nucleobases and RNA backbone. Acceptor to Donor ratio is ∼ 2:1

### Initial exploration of the DBA/ACN PS-DVB

**Figure 2A** shows the charge state envelope of an unmodified RNA 40mer ranging from z=-16 to z=-6 with heavy adduction in the lower charge states. Iribarne and Thompson^24^ showed adduction of a molecule decreases the overall Gibbs Free Energy needed to transition from the liquid phase into the gas phase. For oligonucleotides, adduction is commonly seen with Group 1 alkali metal contaminates (Na^+^, K^+^, etc.) however a closer examination of the mass differences within the spectra here showed that the adduction mass shifts were characteristic of the alkylamine with a smaller peak having a mass difference attributable to an acetonitrile adduct **(Figure 2B)**. The high temperatures used in the source should effectively remove the low boiling point solvent, however, it has been shown that due to its lone pair of electrons, acetonitrile can act as a hydrogen bond acceptor^37,38^, and unlike other common protic solvents used in chromatography this effect results in microheterogeneity of this solvent system.^39^ It is therefore unlikely the acetonitrile adduction occurs at the charged phosphodiester site but instead is occurring on the Watson-Crick face of the oligo by a hydrogen bond donor. Adduction was cleared using In-Source Fragmentation (ISF),^24^ of ∼ 35V [**Supplemental Figure 1**] ISF increases the energy in the system to allow for the “cooling” effect of dissociation;^40^ however, the adduction would persist in the liquid phase to impact chromatographic selectivity and retention.

**Figure 2:**
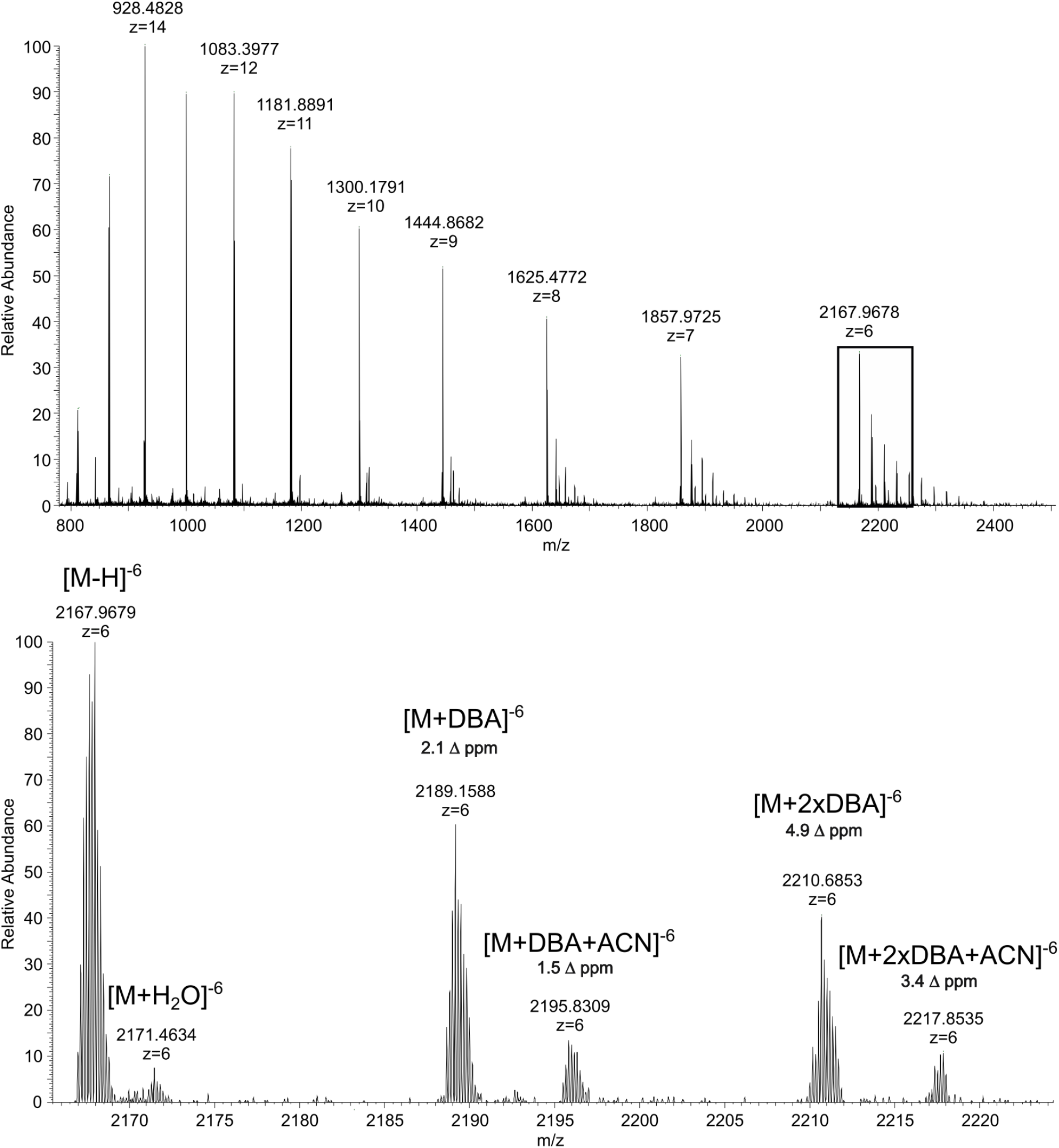
Top Pane: Mass spectra of an RNA 40mer using dibutylamine as the alkylamine. Lower charge states show heavy adduction. Bottom Pane: Zoomed spectra of charge state -6 showing adducts having a mass shift corresponding to the alkylamine and the organic. High resolution accurate mass deconvolution shows mass shift accuracies of less than 5 ppm.

**Figure 3:**
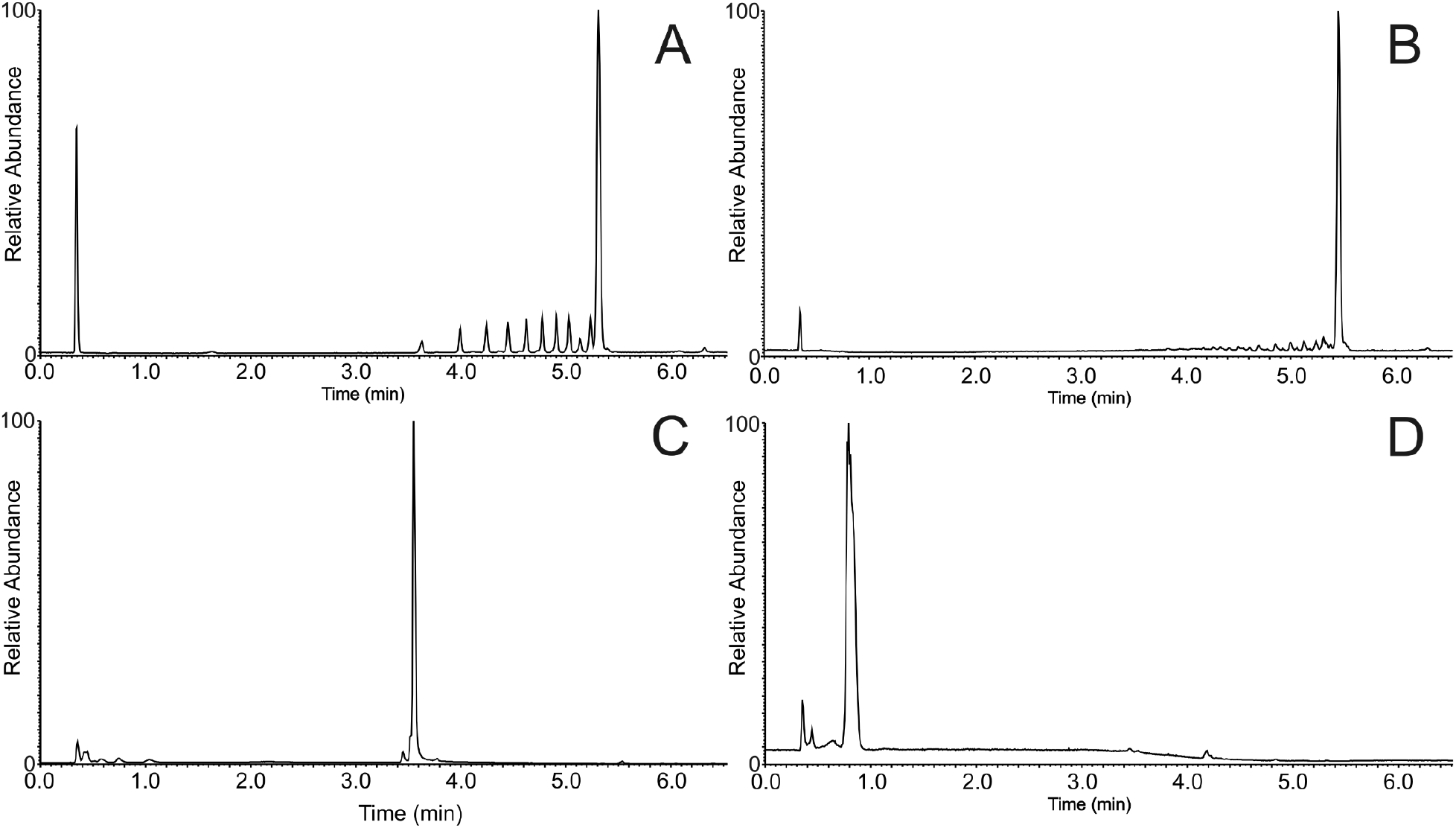
Total ion chromatograms of polydeoxyadenosine and polydeoxyinosine under hydrogen bond donor or acceptor alkylamine. **A**: polydeoxyadenosine with 5 mM DBA **B**: polydeoxyinosine with 5 mM DBA **C**: polydeoxyadenosine 5 mM TEA **D**: polydeoxyinosine 5 mM TEA. Note, smaller peaks in **A** and **B** are failure sequences from synthesis.

### Liquid Chromatography Mass Spectrometer through Hydrogen Bonding

To test the effect of hydrogen bond/donor acceptor mechanism on the nucleobase during ion-pair chromatography, two oligonucleotides were designed. One oligonucleotide consisted of poly deoxy inosine with the second being poly deoxy adenine, both containing 5’ and 3’ hydroxyl groups. This results in the poly-inosine construct having 17 hydrogen bond donating sites, the two terminal hydroxyls and the hydrogen at N1. The poly-adenine construct has the same 17 hydrogen bond donating sites plus the N6 amine, making a minimum of 32 sites, roughly double that of the inosine.

The alkylamine triethylamine (TEA) is a tertiary amine, and can only function as a hydrogen bond acceptor, whereas the secondary amine, dibutylamine can function as both a hydrogen bond donor and acceptor. **Figure 2** shows the results of a comparison of the acceptor amine TEA with that of the donor/acceptor amine DBA on both constructs. In the presence of DBA, retention is observed for both polyA and polyI, however for TEA, retention is only observed with the polyA oligonucleotide. The system’s pH is ∼ 9.0, producing a single negative charge for each phosphodiester linker and deprotonation at position N1 on the purine ring. Both DBA and TEA carry a positive charge at pH 9.0 so retention for either amine should function on the classical mechanism of ion paring along the backbone. However, for TEA we see retention only with the polyA construct, where both polyA and polyI would have the same number of electronegative sites to participate in pairing with TEA. Retention can be explained by the formation of a hydrogen bond between the N6 hydrogen(s) of the adenine base and the alkylamine. For the polyI construct the hydrogen at the N1 position is predominately deprotonated and could participate in dipole-dipole interaction with the tertiary amine TEA, however due to the purine ring structure having delocalized electrons, resonance would lessen the availability of the dipole for interaction with TEA. While the interaction with TEA and the polyI construct is restrained on the Watson-Crick face, electrostatic interaction with TEA and the phosphodiester backbone should still allow for retention, although it is not observed. The lack of retention is explained through restricted interaction between the chromatographic substrate and the ion paired oligonucleotide. The physical difference between the PS-DVB surface and a carbon functionalized silica bead is that the ion paired oligo sees an infinite flat surface for the PS-DVB substrate where the alkyl chain on the silica bead offers structural framework. Intercalation of an alkylamine^18^ with the surface bound alkyl chains would create a surface with multiple points of interactions between the surface bound alkyl chain and the ion paired TEA, where interaction with the planar PS-DVB would be limited.

To test for steric hinderance, we analyzed the polyI construct on a C4 and C8 substrate under low and high pH conditions. Alkyl chain substrates on silica supports may contain some level of silanol activity, even when passivated through chemical bridging. To negate any possibility of interaction with residual chemical groups at the surface, the system was adjusted to pH 6 to protonate the substrate and the nucleobase. Figure 4A shows the TEA paired polyI construct retained on C4 media under full protonation. Under these conditions, the oligonucleotide is retained through classical dipole-dipole interaction with the surface intercalated TEA as well as increased hydrophobic interaction by hydrogen bonding of the N1 hydrogen with that of TEA.

**Figure 4:**
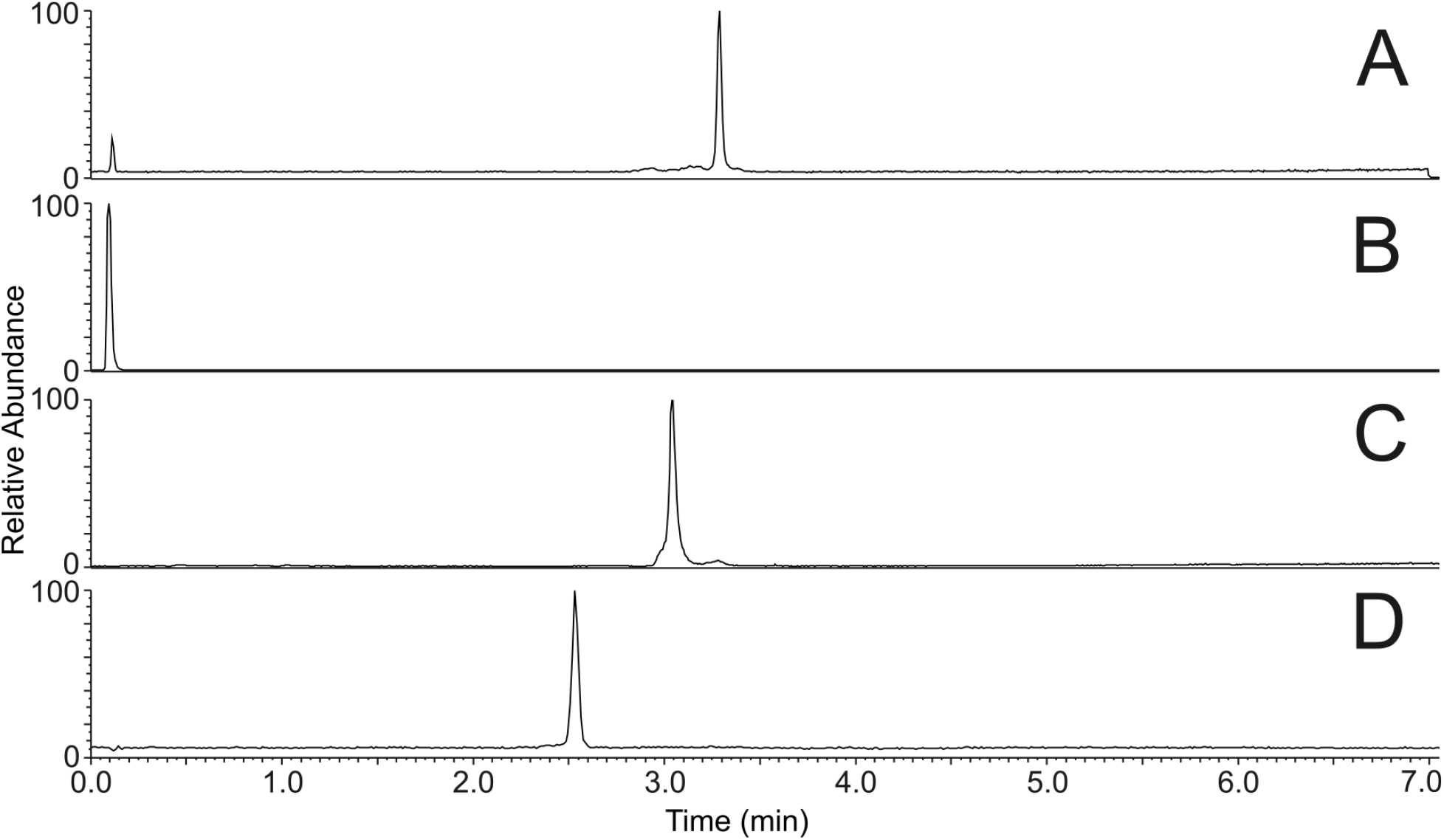
Total ion chromatograms of polydeoxyinosine on C4 and C8 chromatographic media under low and high pH with TEA and DEA as the alkylamine. **A**: Polydeoxyinosine on C4 media pH 6.0 with TEA. **B**: Polydeoxyinosine on C4 media pH 10 with TEA. **C**: Polydeoxyinosine on C8 media pH 10 with TEA. **D**: Polydeoxyinosine on C4 media pH 10 with DEA.

**Figure 5:**
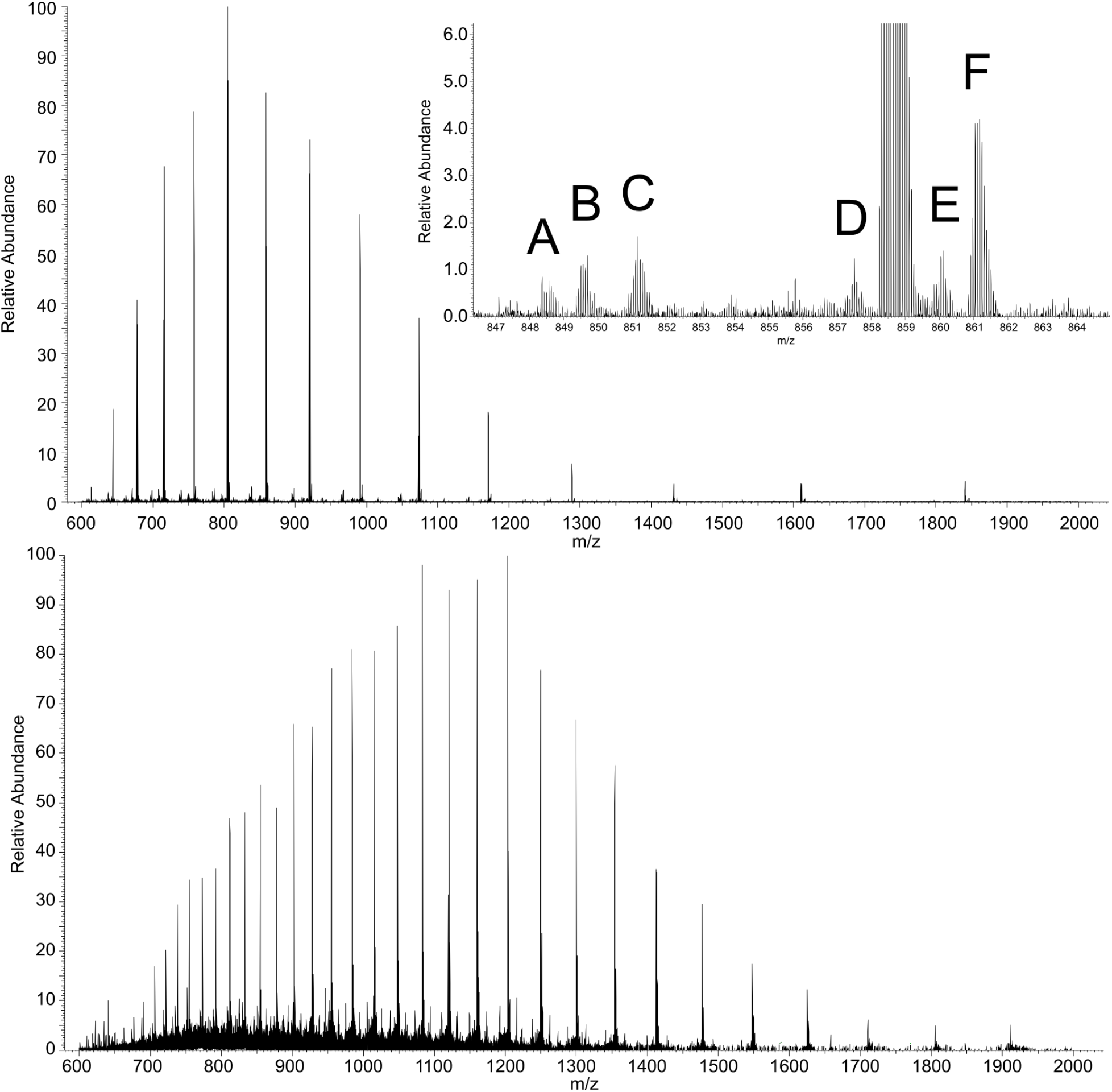
LCMS chromatogram of a 40-mer, top, and a 100-mer, bottom, analyzed on a PS-DVB column with 5mM DBA and “no HFIP”. Inset shows zoomed baseline of highest charge state with assigned impurities detected at 1% relative abundance. **A**: Guanosine nucleotide loss **B**: Adenosine nucleotide loss **C**: Cytidine nucleotide loss **D**: Cytosine loss **E**: Sodium adduct **F**: Potassium adduct

When the pH is adjusted to 10, the N1 hydrogen is no longer available to participate in hydrogen bonding with TEA. Instead, the site becomes available for binding with water, which decreases the number of points of interaction between the oligonucleotide and the chromatographic substrate, resulting in no retention. (Figure 4B). When the chromatographic substrate is changed to C8, retention at pH 10 is observed. Here, the alkyl chain of the C8 offers an increased number of points for dipole interaction with the phosphate backbone than that of C4 due to the increased flexibility of the longer alkyl chain^41,42^ (Figure 4C). Furthermore, interaction with any possible surface silanol activity is reduced. When the alkylamine is substituted with diethylamine, a hydrogen bond donor amine of similar size, retention on the C4 substrate at the elevated pH is observed (Figure 4D).

### Characterization of sgRNA by Intact Mass Analysis

A secondary observation with the use of DBA is minimal adducts detected in the spectra. When examining the spectra of TEA we see common metal adduct Na+ K+, etc. However, in the DBA spectra these adducts are minimal or absent (**Supplemental Figure 2**). Should the mechanism for ionization in the DBA analysis be charge residue, we would expect to see the metal adduction at a high abundance. The lack of metal adducts, the presence of acetonitrile adducts, and the need for source fragmentation to clear the spectra all indicate that the ionization method is through an ejection mechanism.^25^ While molecular dynamic simulations can be used to calculated the change in enthalpy, it is probable that the Gibb’s free energy needed to transition to the gas phase from the electrosprayed droplet is decreased by making the oligonucleotide more hydrophobic through the hydrogen bonding of the alkylamine to sites on the oligonucleotide other than the phosphodiester backbone. Furthermore, the use of fluoroalcohols was initially added to the mobile phase to aid in desolvation by reducing the surface tension of the electrosprayed droplet.^28^ If an oligonucleotide can transition more efficiently through increased hydrophobicity of hydrogen bonded amines, then the need for the fluoroalcohol is less important to the analysis. We sampled an unmodified RNA 40mer and 100mer using a 5 mM DBA mobile phase with the PS-DVB column without the fluoroalcohol added to the mobile phase. **Figure 6A** shows the mass spectrum of the 40mer with a charge state envelope of –7 to –21. The inset shows the zoomed baseline image of the most intense charge state (-16). A white paper by Capaldi, et al. suggested that impurity reporting limits from oligonucleotides be commensurate with the size of the oligonucleotides.^43^ More recently the FDA proposed “phase appropriate” purity determinations,^44^ due to the size of sgRNA used in gene therapies a reporting threshold of 0.5 - 1% is typically adequate for early phase development of gene therapy products. Here we are able to assign impurities in the 40mer strand at 1% relative abundance and within 3 ppm mass error. Terminal nucleotide loss and metal adducts are the most prevalent in the spectra. For the 100mer, baseline resolving impurities at 1% is difficult due to spectral interference arising from signal decay in the orbitrap analyzer.^45,46^ However, impurities are still distinguishable at abundance of ∼ 5% and are verified through isotope simulation using the chemical formula of known adducts (**Supplemental Figure 3**). Both oligonucleotides are easily deconvoluted down to their monoisotopic masses, with delta masses differing by as little as 3 ppm. Given the concerns about PFAS in the environment, removal of HFIP from mobile phases could be a way for some industries to lessen their chemical footprint on the environment.

## Conclusion

In this study we show that LC-MS sampling of RNA is enhanced through hydrogen bonding between hydrogen bond donor amines and the nucleobase. This binding reduces interaction between the oligonucleotide and the hydration shell which allows the RNA molecule a more facile transition from liquid to the gas phase. By using a classical ion pairing approach on alkyl chain modified silica we show that the N1 site on a 15mer polydeoxyinosine construct is occupied by the alkylamine, creating points of interaction between the construct and the chromatographic substrate. Application of this approach with DBA on a polystyrene divinylbenzene substrate resulted in high resolution mass spectra for a Crisper/Cas13 and Crispr/Cas9 generic gRNAs, without the need for fluoroalcohol. Removal of the fluoroalcohol for intact mass characterization of guide RNAs can prove beneficial for labs which are trying to reduce their use of poly fluorinated compounds.

## Supplementary Information

**Supplemental Figure 1:**
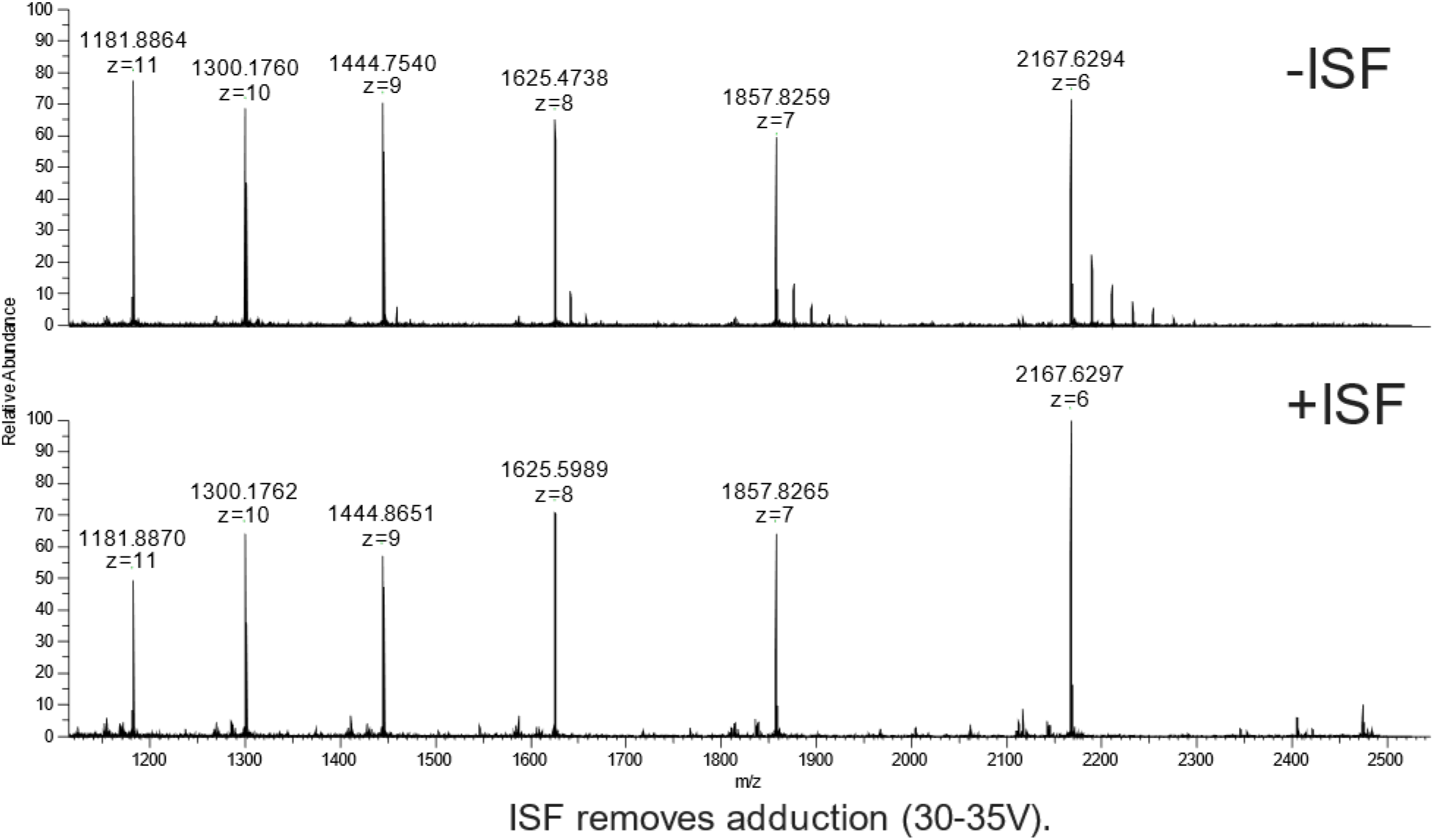
In-Source Fragmentation (ISF) can remove adduction formation prior to MS analysis

**Supplemental Figure 2:**
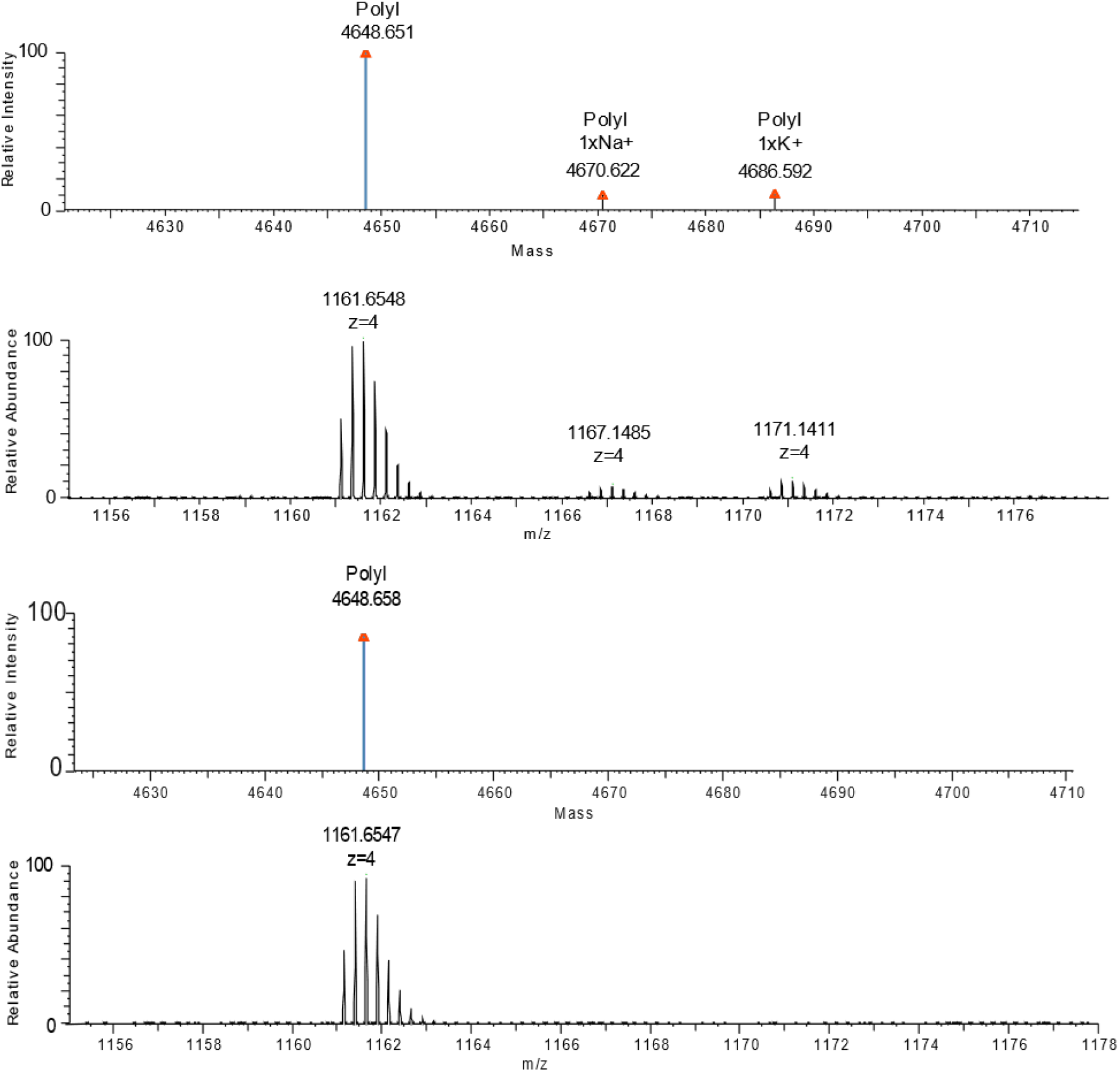
Displacement of metal ions in the presence of DBA

**Supplemental Figure 3:**
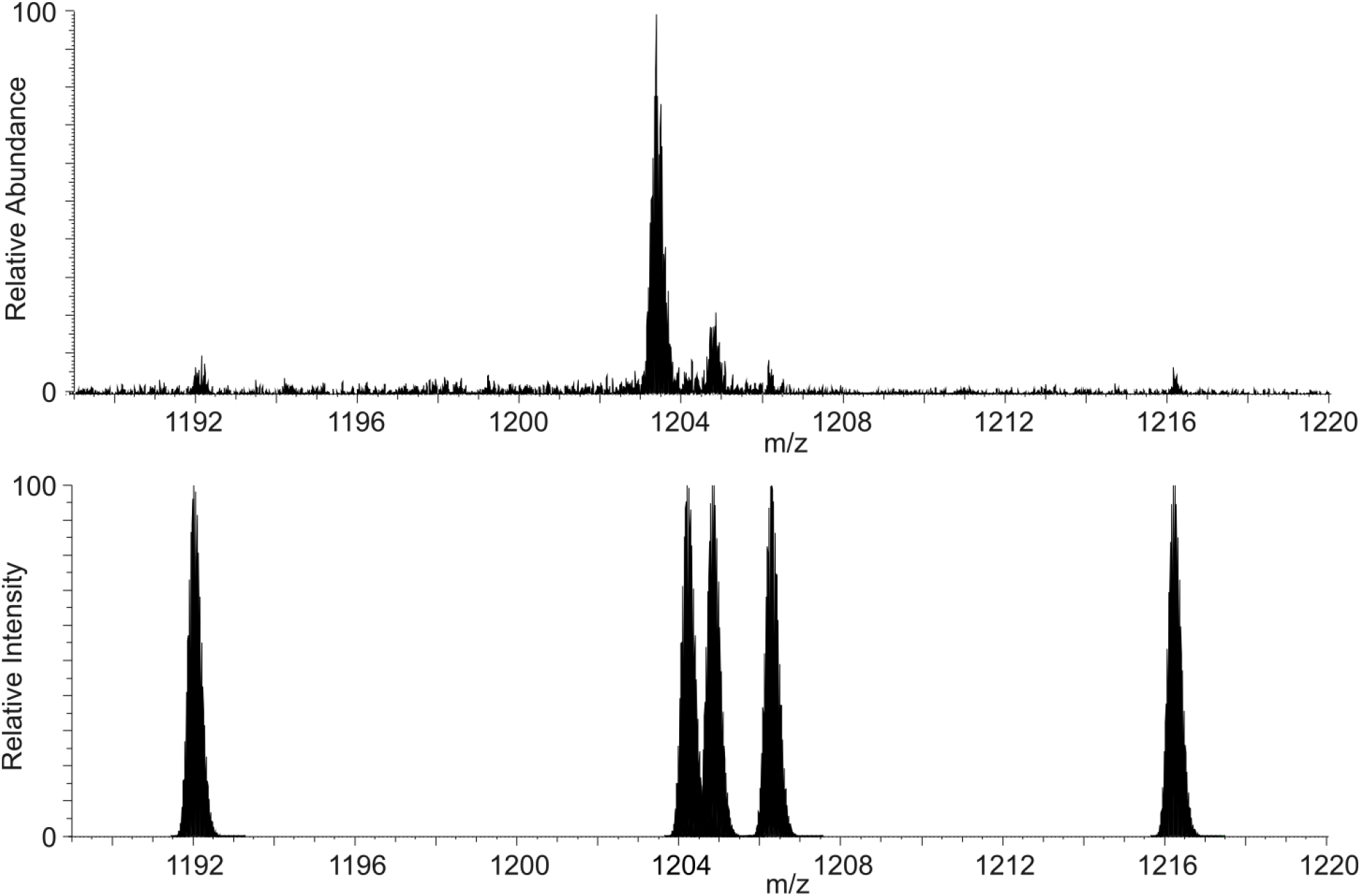
Impurity detection of the 100 gRNA. Tope pane is zoomed baseline of the -27-charge state. Bottom pane shows isotopic simulated peaks for the chemical formulas of known impurities. From left to right: loss of cytidine nucleotide, Na+ adduct, K+ adduct, 2xK+ adduct, guanosine nucleotide addition.

